# Tripartite loops reverse antibiotic resistance

**DOI:** 10.1101/2025.01.04.631305

**Authors:** Farhan R. Chowdhury, Brandon L. Findlay

## Abstract

Antibiotic resistance threatens to undo many of the advancements of modern medicine. A slow antibiotic development pipeline makes it impossible to outpace bacterial evolution, making alternative strategies essential to combat resistance. In this study, we introduce cyclic antibiotic regimens composed of three drugs or “tripartite loops” to contain resistance within a closed drug cycle. We show that as bacteria sequentially evolve resistance to the drugs in a loop, they continually trade their past resistance for fitness gains, reverting back to sensitivity. Through fitness and genomic analyses, we find that tripartite loops guide bacterial strains towards evolutionary paths that mitigate fitness costs and reverse resistance to component drugs in the loops and drive levels of resensitization not achievable through previously suggested pairwise regimens. These findings open the door to sequential antibiotic regimens with high resensitization frequencies which will improve the longevity of existing antibiotics even in the face of antibiotic resistance.

**TOC:** 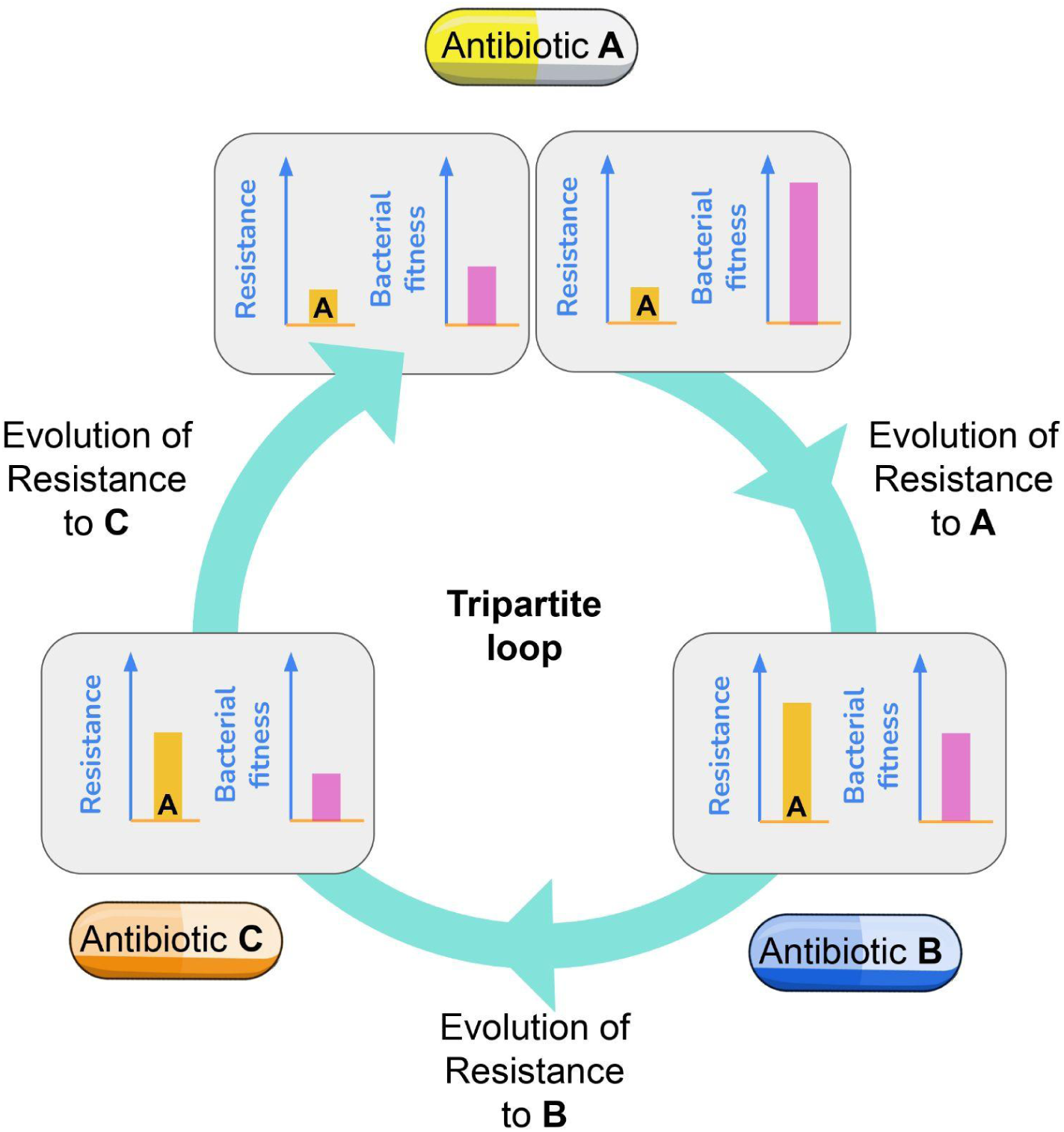

## Introduction

Bacterial infections claim 7.7 million lives each year, of which 4.95 million are associated with antibiotic resistance [1]. The slow pace of antibiotic development is failing to keep up with bacterial evolution, pushing us towards a post-antibiotic era [2–4]. Tipping the scales in our favor in the fight against antibiotic resistance will require alternative strategies beyond the discovery or invention of new drugs to combat antibiotic resistance. One potential approach to slow down resistance evolution is to employ existing drugs in a sequence, with drugs administered one after the other at either predetermined times or as resistance arises [5,6]. Experimental and computational evolution studies indicate that sequential antibiotic regimens can constrain resistance evolution [7–12], and incorporation of collateral sensitivity (CS) is thought to allow maintenance of sensitivity to the alternating drugs indefinitely [10,13–15].

Unfortunately, studies on the effectiveness, importance and repeatability of CS have produced mixed results [16]. Some experimental evolution studies report repeatable CS interactions [9,10,17], while others show weak reproducibility [17–21]. Reports also suggest that sequential antibiotic therapy can constrain resistance evolution independently of CS [21,22]. While the effect of CS on resistance evolution is anchored in several excellent studies which have identified a number of possible drug pairings, most pairings have been experimentally verified using a relatively limited number of evolutionary replicates (2-8 replicates in general) [7,10,15]. Reproducibility is critical for the use of CS in the clinic, and the evolutionary trade-offs that are at the core of sequential or cyclic regimens must be repeatable. Absent large scale experimental evolution studies it is unclear which drug cycles will fail, how often they will fail, and whether those failure rates can be limited.

In a previous study [23], we showed that in a gentamicin (GEN) - piperacillin (PIP) pairwise cycle previously suggested for cyclic therapies [10,15], GEN resistant *Esherichia coli* lineages frequently evolved hypersensitivity towards piperacillin (PIP) but subsequent PIP evolution failed to reverse resistance or reduce adaptation rates, predominantly producing multidrug resistant mutants instead. The repeatability of CS evolution was low even in some previously reported CS-pairs, and there was a lack of complete antibiotic resensitization in most of the pairs tested [23]. This showed that CS interactions often fall apart due to lack of repeatability of evolution, and that pairwise cycles often do not achieve the level of resensitization required for cyclic regimens.

In this study, we ask if mutants that fail to be resensitized in a pairwise cycle can be salvaged, with susceptibility to one or both of the initial antibiotics restored. Although resistant mutants possess strong selective advantages in environments containing the antibiotic of interest, those mutations render them less fit in antibiotic free environments [24,25]. Reversion to susceptibility is then favoured, either through outcompetition by naive cells or by compensatory mutations that enhance fitness but lower resistance levels (phenotypic reversion) [22,26,27]. As it is infeasible to prescribe an antibiotic-free period during an ongoing infection, we instead incorporate a third antibiotic into the series, creating a tripartite loop (Figure 1A). We choose this third drug with a mechanism of action distinct from the other two, limiting the potential for cross resistance [28,29]. We evolve replicates of *Escherichia coli* K-12 substr. BW25113 (wildtype, WT; n = 16) through experimental tripartite loops using a soft agar gradient evolution (SAGE) based platform [30]. The large sample size allows us to capture repeatable evolutionary outcomes, and our experimental design allows identification of evolutionary trade-offs that are robust against compensatory evolution [12,16,23,24]. We find that evolution of nitrofurantoin (NIT) resistance reliably restores GEN susceptibility in bacteria resistant to GEN and PIP when bacteria are evolved against drugs in the order GEN-PIP-NIT. This loop is effectively bidirectional, with NIT resistant bacteria reliably resensitized through a NIT-PIP-GEN loop. This effect is not limited to NIT, as a suboptimal drug like doxycycline (DOX), against which the majority of the GEN and PIP-resistant strains were cross-resistant, was able to reinstate GEN sensitivity. We find that to bypass the fitness loss associated with multidrug resistance, cells rewire their metabolic pathways, concurrently restoring susceptibility to the first drug in the series. All resensitizations we observe occur independently of CS interactions between component drugs in the loop. Overall, we demonstrate that in some cases the multidrug resistance that arises in pairwise loops can be reversed by extending to tripartite loops, experimentally validating a path to more effective and more resilient cyclic antibiotic therapies.

**Figure 1:**
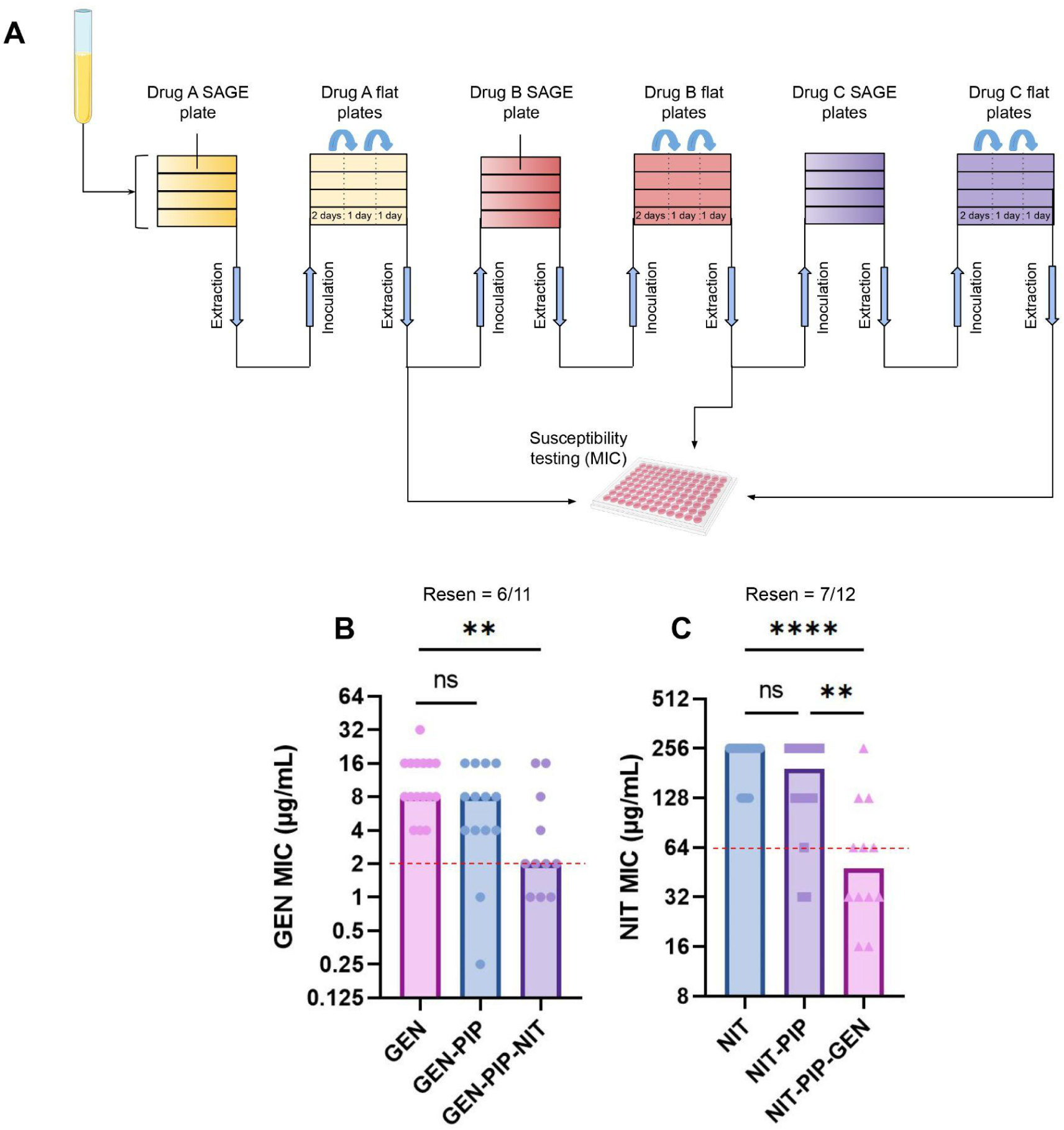
Tripartite loops improve antibiotic resensitization. (A) SAGE is used to study three-drug cyclic regimens or tripartite loops. Bacteria were inoculated into soft agar containing antibiotic gradients to generate resistant mutants (n = 16). SAGE plates were incubated for a fixed duration of 7 days, after which mutants were harvested and passed through three “flat plates” containing the same antibiotic from the prior SAGE plate at a concentration = ½ the evolved MIC of the mutants. The incubation period for each flat passage is noted in the figure. MIC and CS profiles of mutants were determined after the end of the flat plates. (B) GEN MICs of strains that passed through the GEN-PIP-NIT tripartite loop. The y-axis denotes the GEN MICs and the x-axis denotes the sequence of antibiotics against which the strains were evolved. For example, the GEN-PIP bar shows the GEN MICs of strains that were sequentially evolved to GEN and PIP (as shown in (A)). (C) NIT MICs of strains that passed through the NIT-PIP-GEN loop. Resen = resensitization counts. Dotted red lines indicate the clinical breakpoint (EUCAST). Bars represent the median MICs. **p<0.01, ****p<0.0001, Kruskal-Wallis with uncorrected Dunn’s test.

## Results

### Tripartite drug loops that resensitize bacteria to antibiotics

We previously reported using soft agar gradient evolution (SAGE) to generate 16 independent replicates of *Escherichia coli* K-12 substr. BW25113 (WT) resistant to both GEN and PIP [23], a drug pair previously proposed to promote resensitization [10,15]. Out of the 16 strains,two strains went extinct during PIP evolution, while the majority of the remaining 14 maintained resistance to GEN (Figure 1B) (Supplementary Figure 1A, F) [23]. In this study, we screened for drugs that could resensitize these strains to GEN, extending our experimental design to incorporate evolution against a third drug “C” (Figure 1A). SAGE plates generated antibiotic resistant mutants, while flat plates containing sub-inhibitory concentration of the challenge antibiotic allowed compensatory evolution to generate fitter mutants [23,24].

Evolution of NIT resistance reduced the GEN resistance of seven out of eleven strains to or below the clinical breakpoint, while driving three lineages extinct (Supplementary Figure 1A), with a median 8-fold drop in GEN MIC (Figure 1B, Supplementary Figure 1F). To account for possible random fluctuations in MIC measurements [31] affecting resensitization counts, we defined antibiotic resensitization as a four-fold or greater reduction in MIC compared to levels evolved when they first encountered the antibiotic, in addition to reduction at or below the clinical breakpoint. Using this definition, NIT resistance resensitized six out of eleven strains to GEN (Figure 1B).

To determine the effect of subsequent evolution against GEN, we subjected the six strains to GEN SAGE plates again, keeping the concentration of GEN equal to their first exposure. Although resensitized, the GEN MIC of these strains were 2-4-folds higher than the WT, making this a 2-4-fold smaller GEN challenge than the one faced by the WT (Supplementary Figure 1F)., While we achieved a 100% evolution rate following the first GEN SAGE plate with WT bacteria (Supplementary Figure 1A) [23], 3/6 lineages went extinct in this second exposure (Supplementary Figure 1B). This shows that not only were these strains resensitized to GEN, but their ability to develop GEN resistance was also impaired.

When we measured NIT resistance in the three surviving mutants, we observed a 4-fold to 16-fold reduction in NIT resistance levels, rendering all three strains resensitized to NIT (Supplementary Figure 1D). This hinted at a possibility of bidirectionality in this loop, where GEN and NIT resistance were mutually exclusive. To test this at scale, we restarted our evolution experiments with 16 replicates, this time evolving resistance sequentially to NIT, PIP and then GEN. Following evolution against GEN we saw a ∼5-fold reduction in median NIT resistance (Figure 1C). Nine out of 12 strains that completed this challenge fell at or below the NIT resistance breakpoint, with 7/12 reaching resensitization (Figure 1C) (Supplementary Figure 1G). There were no extinctions on exposure to NIT or PIP, but four strains went extinct during GEN evolution (Supplementary Figure 1C).

When strains were evolved sequentially to PIP, GEN and NIT, NIT had no significant impact on PIP susceptibility (Supplementary Figure 1E). This showed that ordering of GEN, PIP and NIT was critical for achieving resensitization, but when applied correctly produced significant resensitizations.

### PIP resistance is important for resensitization

Stratifying results from each step of the GEN-PIP-NIT loop by final GEN MIC revealed that strains which were ultimately resensitized to GEN exhibited decreased GEN resistance following PIP adaptation, while those that maintained GEN resistance after NIT were unchanged by evolution against PIP (Figure 2A, B). Similarly, stratifying NIT resensitized and resistant strains from the NIT-PIP-GEN loop revealed that PIP evolution reduced NIT resistance by 2-fold in the resensitized strains, but not in the resistant ones (Figure 2C, D). Overall, strains evolved through an intervening PIP evolution step exhibited a 4-fold reduction in GEN resistance on NIT exposure, as opposed to a 2-fold difference when the PIP step was omitted (Figure 1B, 2E). This suggests that the incorporation of a third drug allows for resensitizations which would not be possible in pairwise loops.

**Figure 2:**
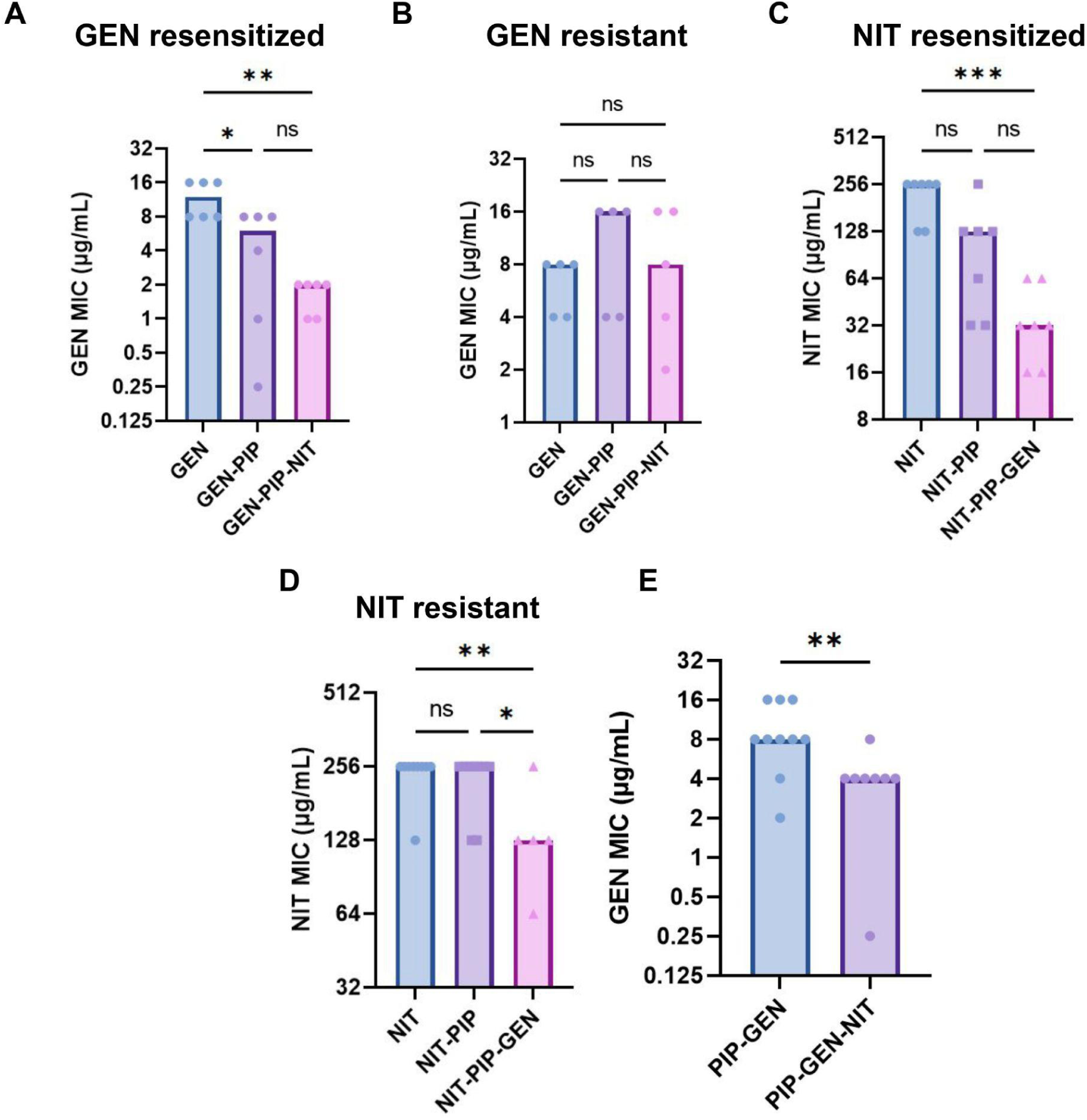
In tripartite loops PIP aids resensitization. (A) GEN MICs of GEN-resensitized and (B) GEN-resistant strains that passed through the GEN-PIP-NIT loop. (C) NIT MICs of NIT resensitized and (D) resistant strains that passed through the NIT-PIP-GEN loop. *p<0.05, **p<0.01, ***<p<0.001, ****p<0.0001, Kruskal-Wallis with uncorrected Dunn’s test. (E) GEN MIC of strains that passed through a PIP-GEN-NIT tripartite loop. MICs after the PIP step are not shown. For all graphs, The y-axis denotes the MICs and the x-axis denotes the sequence of antibiotics against which the strains were evolved before measuring the MICs. For example, the PIP-GEN-NIT bar shows the GEN MICs of strains that were sequentially evolved to PIP, GEN and NIT.**p<0.01, Mann Whitney test. Bars represent the median MICs.

### Resensitizations are independent of CS and principally mitigate fitness loss

To identify the driver of GEN resensitizations in the GEN-PIP-NIT loop, we first examined the effect of forward CS [23] to NIT. To avoid missing even a weak connection between CS and resensitizations we included all strains that showed any reduction in GEN resistance upon NIT evolution in the analysis.

We found no correlation between NIT CS in the GEN and PIP multidrug-resistant strains and reductions in GEN resistance (Figure 3A, B: left column; Supplementary Figure 1F: GEN MICs panel). To test if backward CS [23] helped resensitize bacteria to GEN, we evolved 16 WT strains to NIT (flat plates included) and measured their GEN CS. Only 6/16 of these strains showed 2-fold CS to GEN (and just one with 2-fold PIP CS) (Figure 3B; Supplementary Figure 1G: GEN MICs panel). In contrast, >50% of the strains were resensitized to GEN in the GEN-PIP-NIT loop, with a median 4-fold drop in resistance (Figure 1A). This remained true for the NIT-PIP-GEN loop, with few CS interactions between the drugs (Supplementary Figure 1G). The resensitizations we observed appeared to be largely independent of forward CS, and while backward CS may have played a role, it was not strong enough to resensitize strains to the extent that we observed.

**Figure 3.**
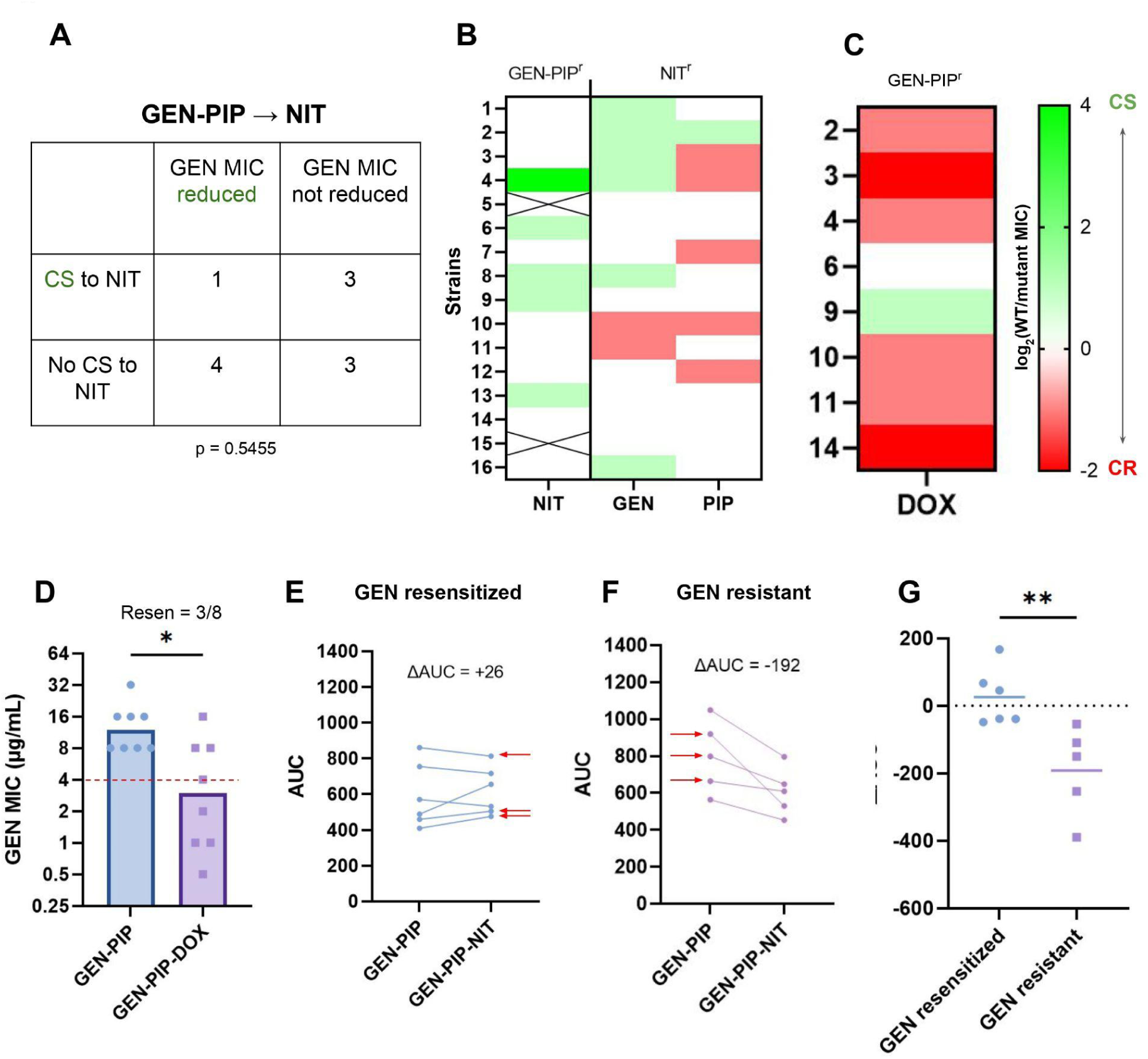
CS does not correlate with resensitizations, but mitigates fitness loss. (A) Contingency table for the 11 strains which evolved NIT resistance through the GEN-PIP-NIT loop, showing no associations between forward CS and GEN resensitizations. Fisher’s exact test. (B) First column: Forward NIT CS of the GEN and PIP evolved mutants from the GEN-PIP-NIT loop. Second column: Backward GEN and PIP CS of WT bacteria evolved to NIT. CS interactions are reported on a log2 scale. (C) DOX MICs of an eight strain subset of the GEN and PIP evolved mutants from the GEN-PIP-NIT loop. CS interactions are reported on a log2 scale. The y-axis denotes the ID of the strains that were picked for DOX MIC testing. (D) GEN MICs of the subset that passed through the GEN-PIP-DOX loop. The x-axis denotes the sequence of antibiotics against which the strains were evolved before measuring GEN MICs. For example, the GEN-PIP-DOX bar shows the GEN MICs of strains that were sequentially evolved to GEN, PIP and DOX. Dotted red line indicate the clinical breakpoint. Bars represent the median MICs. *p<0.05, Mann Whitney test. (E) and (F) AUCs of strains before and after NIT evolution for GEN-resensitized and GEN-resistant strains respectively. The x-axis denotes the sequence of antibiotics against which the strains were evolved before measuring AUCs. GEN-PIP = before NIT evolution, GEN-PIP-NIT = after NIT evolution. ΔAUC is the average of the difference between post and pre NIT AUCs. For the GEN resistant group, we considered every strain that did not meet our resensitization criteria as resistant. This resulted in the inclusion of one strain that was below the GEN resistant breakpoint but did not reach our resensitization standard. Red arrows indicate the strains that were sequenced. (G) ΔAUC of individual strains plotted, grouped by resensitized and resistant. Horizontal lines represent the mean. **p<0.01, unpaired t test.

To test if the specific mechanism that conferred NIT resistance drove GEN resensitization, we evolved GEN-PIP multidrug-resistant lineages against doxycycline (DOX), a tetracycline antibiotic with a different mechanism of action from NIT, GEN, or PIP [32] (n = 8). Despite most of the eight mutants showing cross-resistance to doxycycline (Figure 3C), 5/8 strains dropped their GEN resistance to or below the resistance breakpoint, with 3/8 reaching resensitization (Figure 3D). This provided further support that CS is not required for resensitization, and indicated factors other than specific resistance pathways contributed to the resensitizations we observed.

Next, we hypothesized that the cumulative fitness costs of maintaining multiple drug resistance may promote the adoption of evolutionary paths that ameliorate these costs, resulting in phenotypic reversion. To test this, we measured strain fitness after each evolution step in the GEN-PIP-NIT, NIT-PIP-GEN and PIP-GEN-NIT tripartite loops, using area under growth curves (AUC) as a proxy for fitness (Supplementary Figure 2) [21,24]. In the GEN-PIP-NIT loop we found only a small drop in average fitness after each evolution step, which did not reach statistical significance (Supplementary Figure 2A). However, stratifying strains on the basis of resensitization to GEN revealed a clear difference in fitness (Figure 3E-G). Strains that were resensitized to GEN upon NIT evolution either saw small gains or marginal losses in fitness (Figure 3E, G), while those that retained GEN resistance lost significantly more fitness on average, with none gaining fitness (Figure 3F, G).

The results of the NIT-PIP-GEN loop were less clear. We observed large fitness losses after every step of evolution, with the evolution of GEN resistance in particular producing extremely unfit mutants (Supplementary Figure 2B). Two of the seven NIT resensitized strains exhibited moderate to large fitness gains upon GEN evolution but none of the five NIT resistant strains did (Supplementary Figure 2D-F). However, the differences in ΔAUC between the resensitized and resistant groups did not reach statistical significance (Supplementary Figure 2F).

Strains from the PIP-GEN-NIT loop showed a large fitness drop as they moved from PIP to GEN, but did not show a significant change in fitness following NIT evolution (Supplementary Figure 2C). Given the increased burden of resistance to three separate antibiotics, we expected that a constant AUC would correspond to significant resensitization. However, only one lineage exhibited increased PIP susceptibility, and that was following GEN evolution, not NIT (Supplementary Figure 1E). Looking more closely into the MIC profiles of these strains revealed that while there were no significant changes in PIP susceptibility, 7/8 strains had increased GEN susceptibility following NIT exposure (Figure 3C). As GEN resistance was consistently associated with the largest fitness penalties (Supplementary Figure 2), this may have off-set a fitness penalty from acquiring NIT resistance, while leaving the less costly PIP resistance unchanged. Overall, the tripartite loops led to higher fitness costs and increased resensitization compared to pairwise loops.

### Whole genome sequencing sheds light on resistance and resensitization mechanisms

To understand the genetic basis of the resistance and resensitization observed, we sequenced the genome of six lineages from the GEN-PIP-NIT loop: three that were resensitized to GEN and three that remained resistant to GEN after NIT evolution (Supplementary Figure 1D). Each lineage was sequenced following every evolution experiment (Figure 1A), allowing us to reconstruct all six evolutionary trajectories (Figure 4A-D).

**Figure 4.**
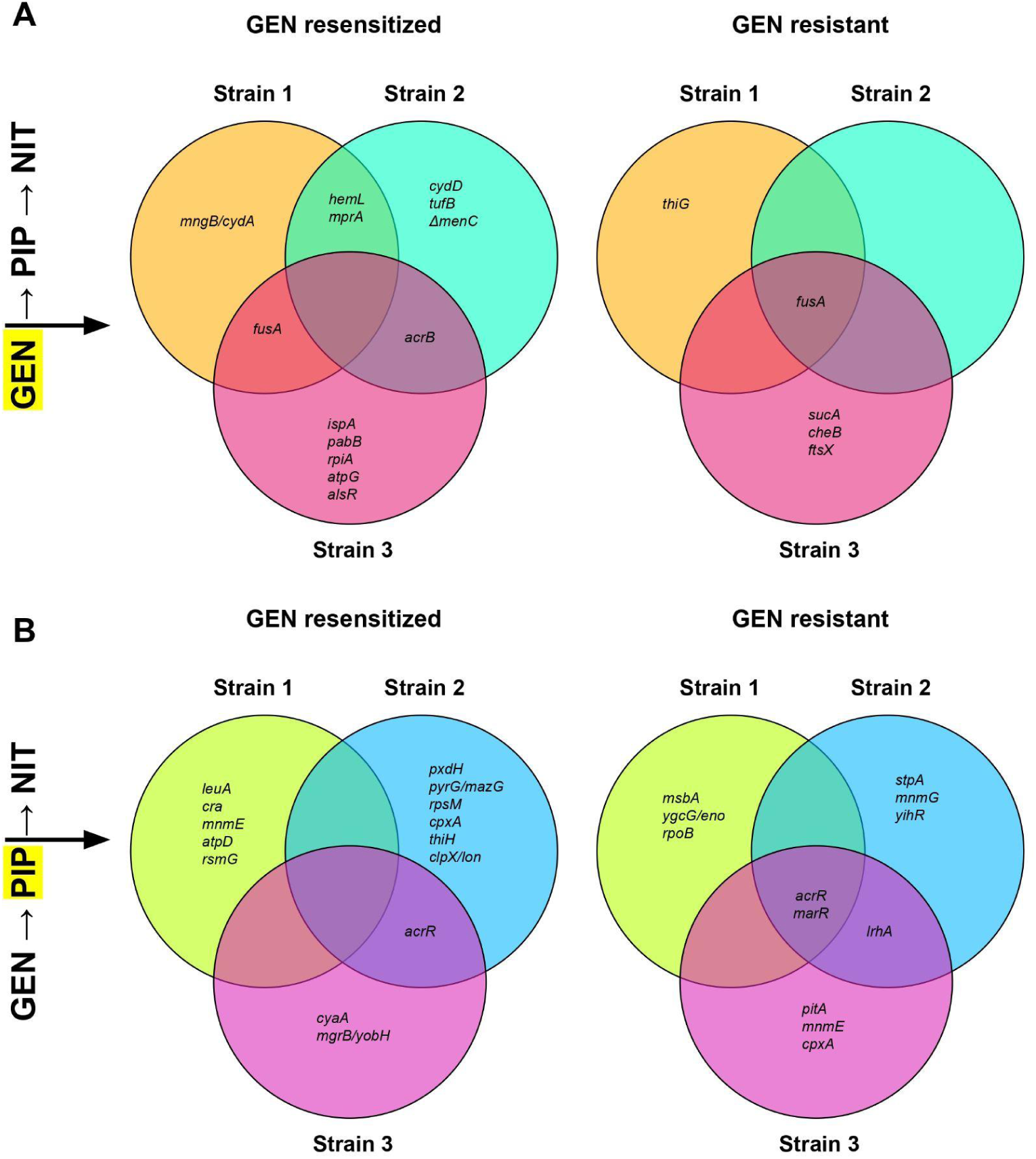
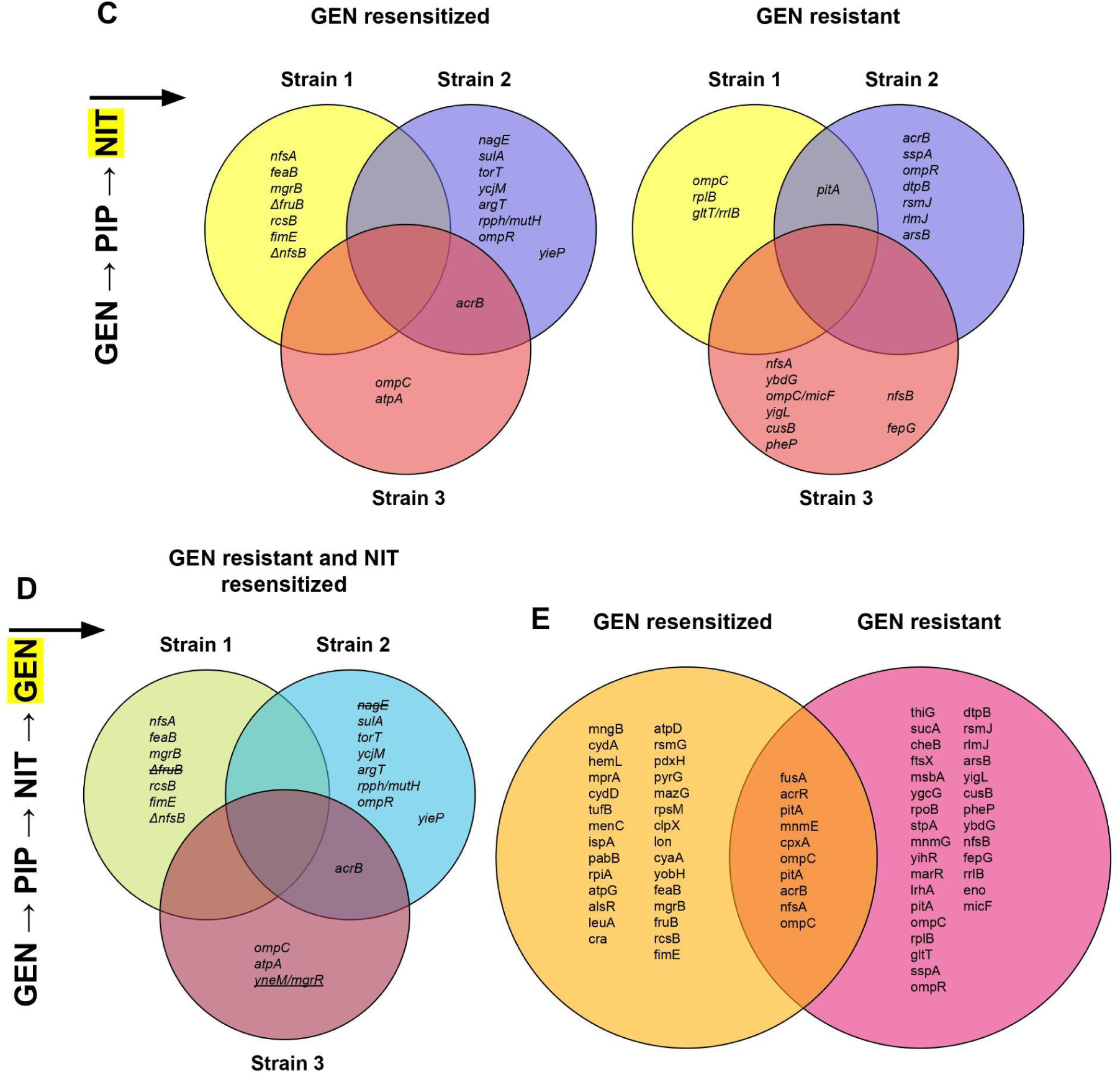
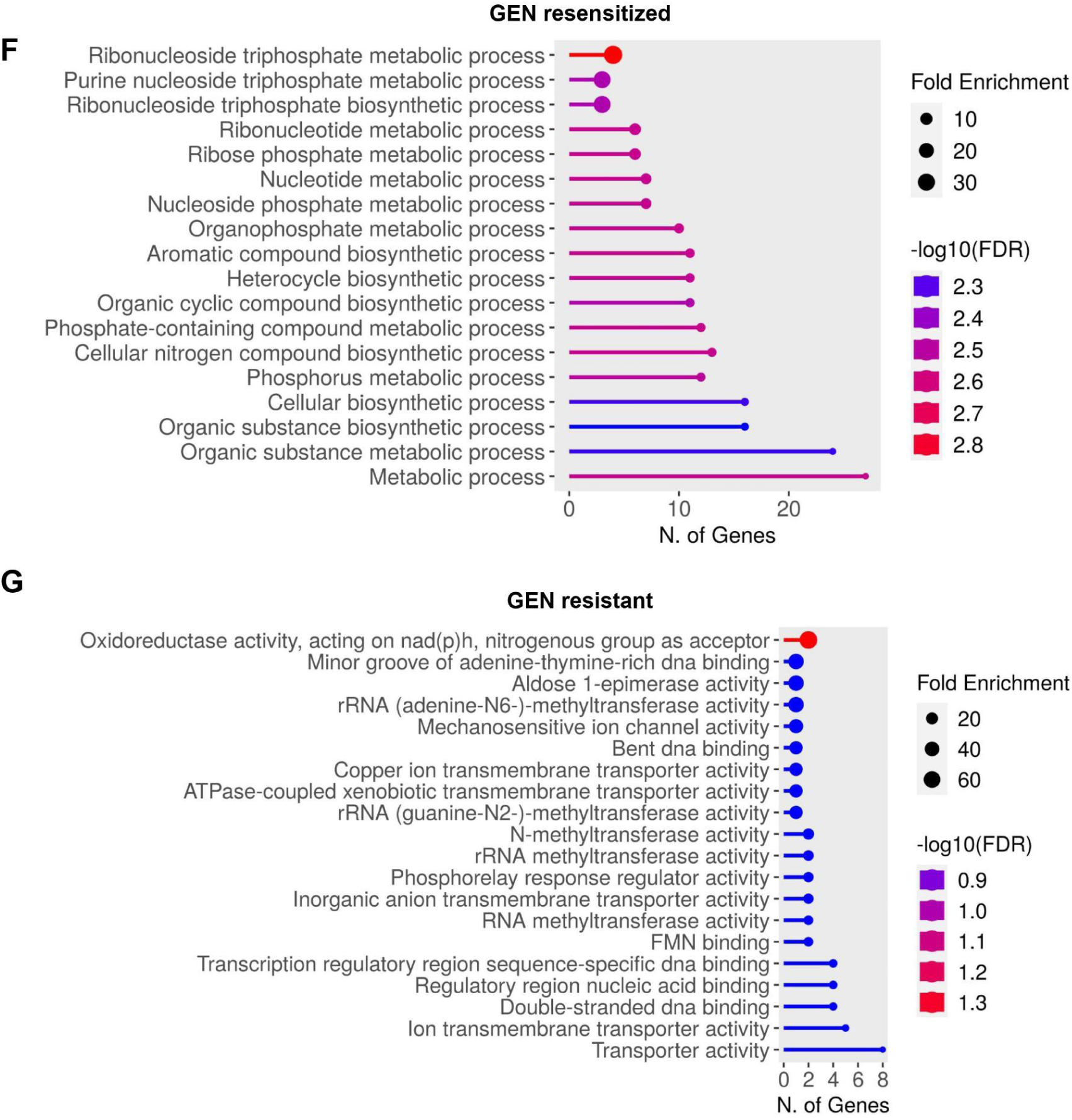
Tracking genomic changes through the GEN-PIP-NIT loop. (A) - (D) Venn diagrams show overlapping and unique mutations in the GEN-resensitized and GEN-resistant strains from the three strains sequenced. The label on the left denotes when the strains were sequenced, with the most recent evolution step highlighted. Strikethroughs denote mutations that appeared in a prior step but were not present in the current step. Underlined mutation in D, strain 3 represents a newly acquired mutation absent from strain 3 in C. (E) Venn diagram showing all overlapping and unique mutations between the GEN resensitized and GEN resistant group, pooled from every step (GEN-PIP-NIT only). (F) and (G). GO term enrichment analysis of unique mutations in the GEN resensitized and GEN resistant groups.

Five of the six lineages acquired their initial GEN resistance through mutations in the translation elongation factor G, *fusA,* mutations that are known to reduce gentamicin’s ability to bind to the ribosome (Figure 4A) [33]. Even though both the GEN resensitized and resistant groups evolved similar GEN MICs (Supplementary Figure 1F), the resensitized strains contained multiple additional mutations in genes involved in the electron transport: *hemL* [34], *cydA* [35], *cydD* [36], *menC* [37], and *atpG* [38] (Figure 4A). Mutations in the electron transport chain can provide GEN resistance either by disrupting drug uptake or reducing ribosomal protein levels [39,40].

Five out of the six lineages acquired mutations in the efflux regulators *acrR* and *marR* following exposure to PIP, changes known to confer β-lactam resistance (Figure 4B) [41]. Mutations in the resensitized group also included other genes involved in β-lactam resistance such as *cpxA* and *cyaA* [42,43]; genes involved in carbon, amino acid, and vitamin metabolism: *cra* [44], *leuA* [45], *pdxH* and *thiH* [46,47]; and the ribosomal genes *rsmG* and *rpsM* [48,49] (Figure 4B). The resistant group did not show any clear mutations in genes involved in metabolism or the ribosome.

All NIT-resistant mutants acquired mutations in one or more of the genes involved in NIT resistance: *nfsA, nfsB* [50], *sulA* (essential for NIT resistance in *lon* mutants) [51], ompR [52] and ompC [53,54] (Figure 4A). Both GEN-resensitized and GEN-resistant lineages showed multiple mutations involved in transmembrane transporters. The resensitized group acquired mutations in genes involved in the sugar phosphotransferase transport system: *fruB* [55] and *nagE* [56], which also have putative roles in aminoglycoside uptake [57], while the resistant strains gained mutations in metal ion, amino acid and peptide transporters instead: *cusB* [58]*, fepG* [59]*, pitA* [60]*, dptB* [61], and *pheP* [62] (Figure 4A). A gentamicin uptake assay suggested that these transport related mutations in the GEN resensitized strains may have slightly increased GEN penetration, but the results did not reach statistical significance (Supplementary Figure 3)

To elucidate how differences between the GEN-resensitized and GEN-resistant groups could affect their propensity towards resensitization, we first identified overlapping and unique mutations between the two groups following NIT evolution (Figure 4E). Common mutations were mostly those known to confer resistance to GEN, PIP, or NIT, as discussed above. To categorize the remainder, we ran GO term enrichment analyses on the non-overlapping gene sets. Every hit from the resensitized group that was above the enrichment FDR cutoff was involved in metabolic processes (Figure 4F), whereas no significant enrichment was found in the resistant group. Manually removing the FDR cutoff (by setting it to 0.99) identified processes involved in transport and DNA-binding (Figure 4G). Mutations in metabolic processes are often involved in compensatory evolution to mitigate fitness costs and phenotypic reversion of resistance [63–66], which supports our observation of the little to no loss (but rather a slight increase) in fitness in the GEN resensitized strains (Figure 3C), in contrast to the significant fitness loss in the resistant group (Figure 3D). These genomic and fitness outcomes suggest that cells become resistant to antibiotics using similar mechanisms, but bifurcate at the level of fitness cost compensation. Cells that adopt pathways that help mitigate their fitness losses also reverse their resistance to the earlier drugs, strongly suggesting a correlation between the two phenotypes.

Since we also saw a surprising drop in NIT resistance after reacquisition of GEN resistance from the GEN-PIP-NIT-GEN series in all three non-extinct lineages, (Supplementary Figure 1D), we looked at the genome sequence of these NIT resensitized strains (Figure 4D). After reacquisition of GEN resistance, the genomic profile of the three strains looked almost identical (Figure 4C and D) except that strain 1 was replaced by a mutant with an intact *fruB* gene possibly via elevation of a low frequency mutant in the population, while strain 2 reverted its *nagE* mutation (Figure 4D). Both genes are involved in sugar transport. It is unclear how reversion of these mutations allowed GEN resistance reacquisition. There are no direct reports of nitrofurantoin being transported inside the cell via these transporters, but both *nagE* and *fruB* have been reported to carry other drugs like streptozotocin and fosfomycin [67,68]. Since the *nagE* and *fruB* mutations are the only differences between the GEN-sensitive-NIT-resistant and GEN-resistant-NIT-sensitive strains (Figure 4C and D), it is likely that these mutations play a role in GEN and/or NIT resistance levels.

## Discussion

Cyclic antibiotic therapies have been proposed as a way to combat the rise of antibiotic resistance [7–11]. The success of such regimens is thought to hinge on CS interactions between the component drugs [10,13–15]. However, to date, CS has seen no application in the clinic since its first description in 1952 [69] and since its proposed benefits in cyclic therapies, partly because of the unreliability and rarity of CS [17–20]. In a previous study we showed that CS, when applied in the correct direction during cyclic therapies, can help resensitize bacteria to antibiotics [23]. However, we observed that the repeatability of CS evolution was low even in drug pairs with reported CS interactions, typified by small resistance drops and low resensitization frequencies, which may readily lead to the emergence of multidrug-resistant mutants [23].

In this study, we explored the potential of extending pairwise regimens to longer “tripartite loops”. We found that the tripartite loop GEN-PIP-NIT significantly improved resensitization frequencies as compared to the previously proposed GEN-PIP pairwise loop [10,15,23], going from ∼14% [23] to ∼54% of lineages (Figure 1B). The loop was also invertible, with NIT-PIP-GEN reliably resensitizing bacteria to NIT (Figure 1C). Resensitizations were independent of CS (Figure 3A, B) (Supplementary Figure 1F, G), and did not appear to be driven by NIT-specific resistance mutations. The resensitization was at least partially independent of drug identity, as extending the GEN-PIP loop with DOX, against which the bacteria showed cross-resistance, produced GEN-resensitized strains (Figure 3C, D).

When GEN-resensitized strains from the GEN-PIP-NIT loop were subjected to GEN again, we found that the antibiotic posed a significant evolutionary challenge, with three out of the six strains going extinct during SAGE (Supplementary Figure 1B). We did not observe extinctions when WT bacteria were exposed to GEN (Supplementary Figure 1A), implying that multidrug-resistant bacteria have constrained evolutionary paths that limit further resistance development. In fact, the three mutants that were able to reacquire GEN resistance dropped their NIT resistance in the process, showcasing the difficulty in maintaining multiple resistance mechanisms.

Unlike in the laboratory, rapid drug cycling in patients may not be possible due to pharmacokinetic factors [12], and the resulting longer evolutionary periods can allow for compensatory evolution which can mitigate evolutionary trade-offs like CS [16,18,70]. While this complicates CS-based cyclic therapies, our study shows that compensatory evolution can be leveraged to drive phenotypic reversion of resistance. We tracked fitness of resensitized and resistant bacteria throughout our tripartite loops, demonstrating that the sequential adaptation to three antibiotics increased fitness penalties compared to pairwise loops (Supplementary Figure 2), possibly due to the need to carry multiple independent resistance phenotypes. Strains could overcome this fitness loss through resensitization, e.g. to GEN (Figure 3E, Figure 4), or could persist with poor growth (Figure 3F) [70]. This interplay was also apparent in the PIP-GEN-NIT loop, again through resensitization of GEN (Figure 3D). Since GEN evolution imposed the largest penalties in our experiments, it appears to be ideal for incorporation into drug cycling protocols. Our GO term enrichment analyses also clearly show evidence of metabolic rewiring associated with compensatory evolution [63–66] in resensitized strains that are missing from the resistant ones (Figure 4F, G), and the fitness and genomic analyses taken together suggests a strong association between compensatory evolution and resistance reversion.

In further support of longer cyclic regimens, we showed that despite the fact that PIP evolution failed to produce significant resensitizations (Figure 1B, C) (Supplementary Figure 1 F, G) [23], it aided in bringing down resistance to the initial drug in both the GEN-PIP-NIT loop and NIT-PIP-GEN loops (Figure 2A-D), which turned to full resensitizations after evolution against the last drug in the series. Additionally, tripartite loops continued to drive bacterial extinction (Supplementary Figure 1A - C), reinforcing prior work on sequential regimens [10,23].

When we compared the NIT resistant vs NIT resensitized strains from the GEN-PIP-NIT-GEN sequence (Figure 4D), we discovered that the genomes of the two groups were almost identical, except that the NIT resensitized strains reinstated two sugar transporter mutations. Elucidation of the exact mechanism of NIT resensitization will require further studies, but our data suggests a possible, previously unreported role for sugar transporters in NIT resistance (Figure 4C, D).

Overall, our findings suggest that tripartite loops can improve antibiotic resensitization and allow continuation of antibiotic cycling even if pairwise cycles fail, without being limited by CS requirements. With our antibiotic development pipeline failing to keep up with resistance emergence, such cyclic therapies may prolong the lifespan of our existing antibiotics.

## Materials and Methods

### Bacterial strain and growth conditions

*Escherichia coli* K-12 substr. BW25113 and the evolved lineages were cultured in Muller Hinton (MH) media at 37 °C. Antibiotics were added to the growth media as needed to grow or isolate resistant mutants from SAGE plates.

### SAGE evolutions

Evolutions were performed as previously described [23]. SAGE evolved mutants were extracted from within 1.5 cm of the end of the plates after seven days of incubation into MH broth containing the challenge antibiotic at a concentration = 2x the WT MIC. Strains were considered extinct when they could not be recovered after extraction from within 1.5 cm of the end of the SAGE plates [23]. Mutants that went extinct were given a second chance at evolution using the same parameters as before. This allowed us to maintain a larger sample size through the extinction events that occurred at different steps of evolution, and we report both the initial and final extinction counts (Supplementary Figure 1A - C). Antibiotic concentrations in SAGE are listed below, and were determined from trial SAGE experiments to reliably evolve strains with MICs above the clinical breakpoints for each antibiotic [71].

| Antibiotic | Concentration ($\mu\text{g/mL}$) |
| --- | --- |
| GEN | 5 |
| PIP | 40 |
| NIT | 80 |

### Susceptibility testing

Minimum inhibitory concentrations (MIC) of antibiotics were determined using the broth microdilution method as described by the CLSI [72]. Antibiotics were diluted in MH broth, then serially diluted across 96 well plates. Bacteria were inoculated at a concentration of 1/200 of a 0.5 McFarland standard. Plates were incubated overnight at 37 °C without shaking, and the MIC was recorded as the lowest antibiotic concentration that prevented visible bacterial growth.

### Flat plates

Flat plates were prepared as previously described [23]. First, the evolved MIC of the antibiotic used in the preceding SAGE plates was determined for all strains that completed SAGE evolution. Next, specific lanes were created for each strain by pouring approximately 12 mL medium supplemented with the antibiotic at half the MIC of that strain into four-well dishes. This allowed for the maintenance of the resistance gained from SAGE during compensatory evolution. Each replicate underwent three consecutive passages on these flat plates (Figure 1A). The first plate was incubated for two days, and the second and third plates for one day (Figure 1A). Unlike during SAGE evolutions, where extractions were limited to within the final 1.5 cm of the plates, cells from flat plates were extracted from the farthest point of growth.

### Fitness measurements

Growth curves for each strain were made by tracking absorbance readings at 595 nm of 1/100 dilutions of overnight cultures using a plate reader (Tecan Sunrise) for 24 h. Plate lids were treated with 0.05% Triton X-100 in 20% ethanol to reduce fogging [73]. AUCs were calculated using GraphPad Prism.

### Whole genome sequencing

Genomic DNA was extracted using the Bio Basic genomic DNA kit (Cat. no.: BS624). Sequencing and variant calling were performed by Seqcenter (USA) on an Illumina NextSeq 2000, and demultiplexing, quality control, and adapter trimming were performed with bcl-convert (v3.9.3). Variant calling was performed using Breseq under default settings [74]. NCBI reference sequence CP009273.1 was used for variant calling. Common mutations were identified using custom R scripts and venn diagrams were based on the output of the R package *ggvenn*.

### Term enrichment analysis

To identify pathways affected by the mutations observed, ShinyGO v0.81 (https://bioinformatics.sdstate.edu/go/) was used. For GEN resensitized strains, the following parameters were used: Species: *Escherichia coli* str. K-12 substr. MG1655 STRINGdb; DB: Go Biological processes; FDR: Default of 0.05. Resistant strains produced no result using these parameters. These parameters were modified by removing the FDR cutoff to produce the results shown in Figure 4G. The modified parameters were: DB: GO Molecular Function; FDR: set to 0.99.

### Gentamicin uptake assay

Gentamicin uptake was measured using a modified version of a previously reported protocol [75]. 300 μL of overnight bacterial cultures were transferred into 30 mL of MH broth in conical flasks and incubated at 37 °C with 250 rpm shaking until log phase was reached. The log phase of each strain was estimated from their growth curves. OD at 600 nm was then measured for each strain, and cells were either concentrated or diluted to reach an OD of 0.4. 100 μL of cells were transferred into microcentrifuge tubes, and GEN was added at a concentration of 100 μg/mL. Tubes were allowed to incubate at 37 °C with 1000 rpm shaking on a heat block for 15 minutes. Tubes were then chilled on ice for five minutes, then centrifuged at 12,000 g for two minutes. 5 μL of the supernatants were used to spot WT *E. coli* seeded MH agar plates, and left to air dry. Plates were incubated overnight, then photographed from a fixed distance. Images were analyzed by fitting circles around the inhibition zones and measuring the area in px^2^ using ImageJ [76]. Measurements were taken from 6 independent replicates for each strain. The three resensitized strains that were sequenced (Supplementary Figure 1F) were also used to perform this test.

## Acknowledgments

This study was funded by the Fonds de recherche du Québec – Santé (FRQS) (269182). FRC is supported by the Fonds de recherche du Québec – Santé (FRQS) (B2X). We thank Dr. L. Freeman for helpful discussions.

---

**Supplementary Figure 1:**
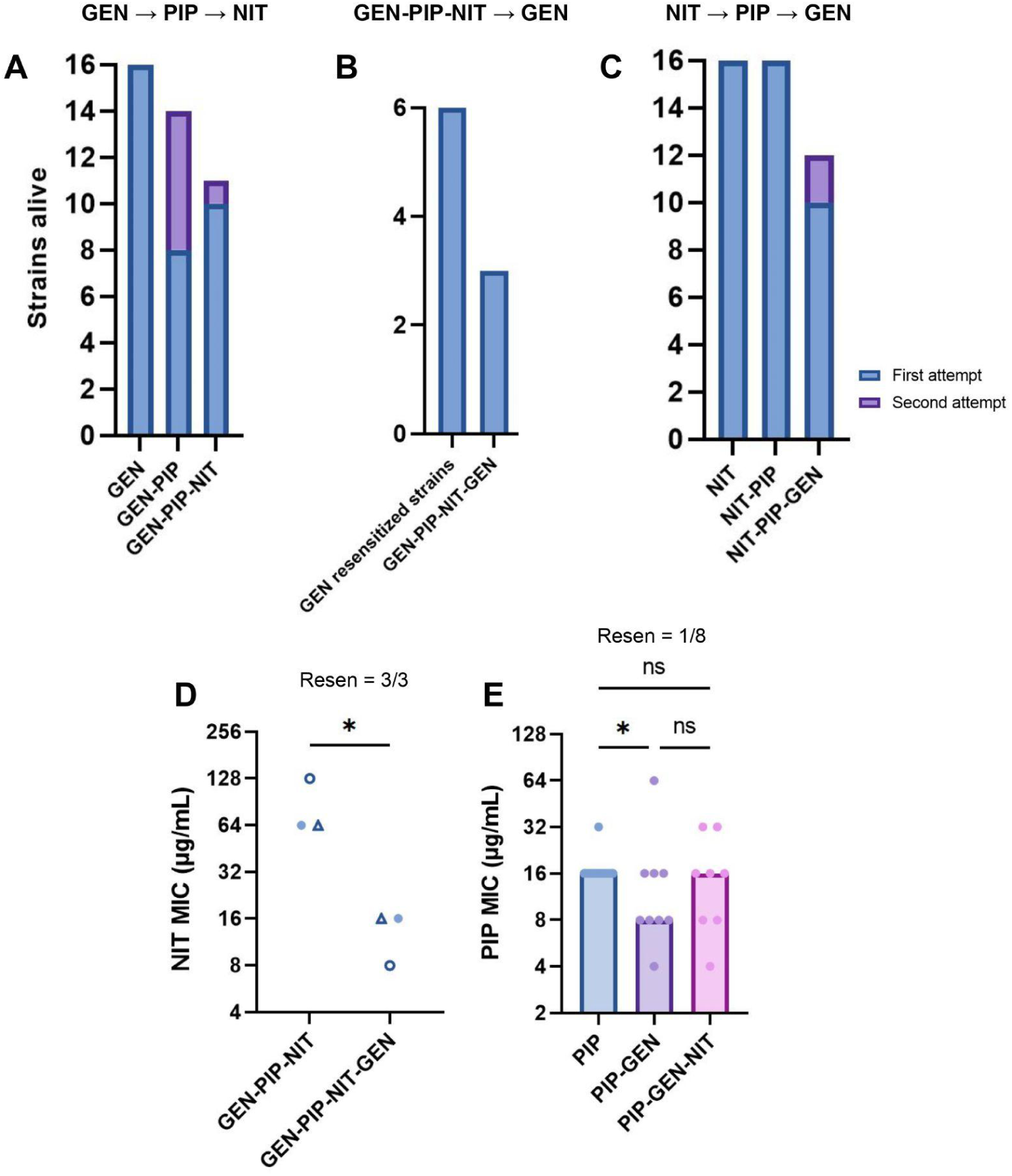
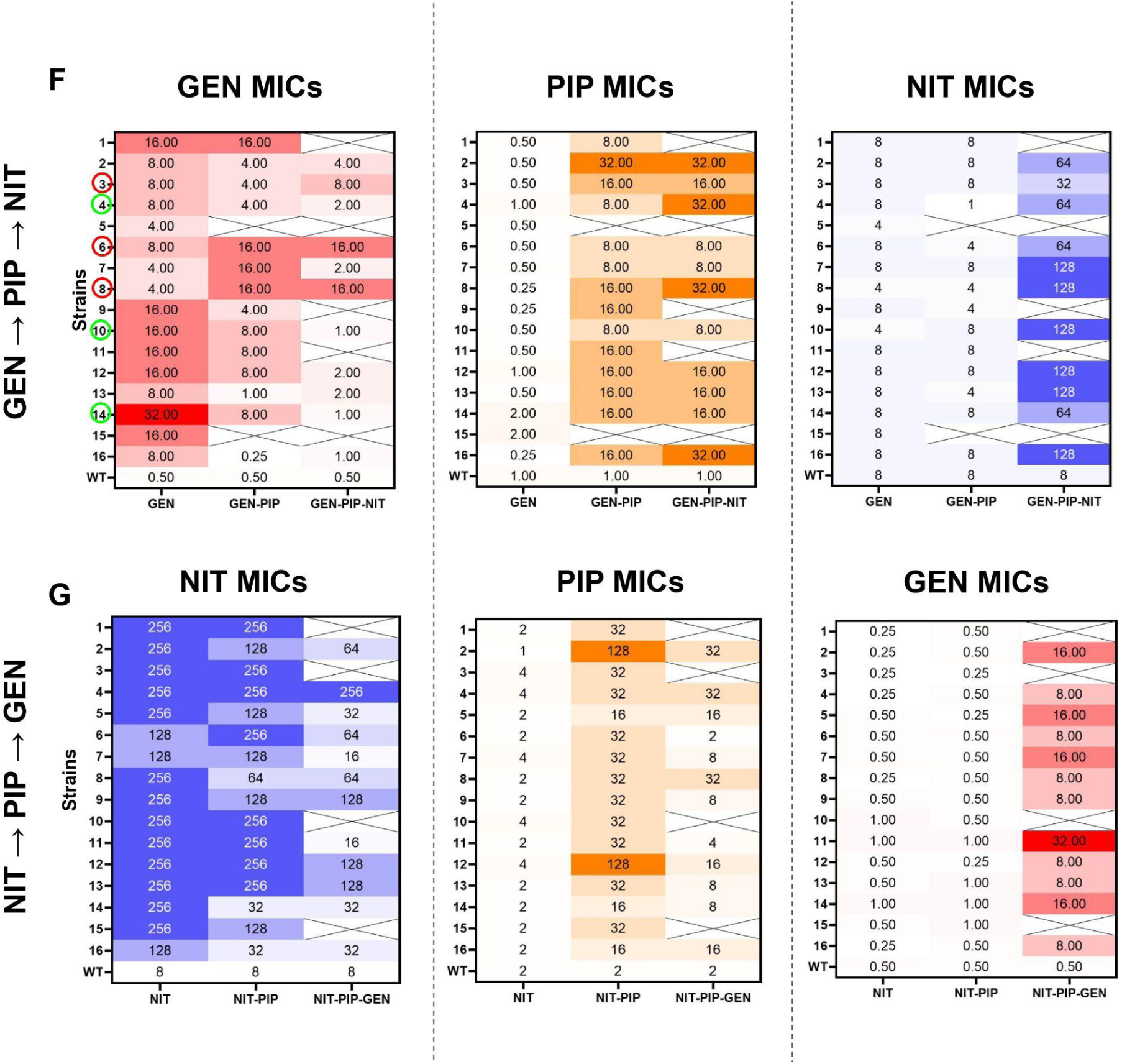
(A) Number of non-extinct strains following each step of the GEN-PIP-NIT loop. (B) Non-extinct strains following the GEN-PIP-NIT-GEN loop. Only strains resensitized to GEN after teh GEN-PIP-NIT loop are included in the analysis. (C) Non-extinct strains following each step of the NIT-PIP-GEN loop. Purple stacked bars denote strains that went extinct on the initial pass, but survived a second attempt. (D) NIT MICs of the three non-extinct after the GEN-PIP-NIT-GEN sequence. *p<0.05, unpaired t-test (E) PIP MICs of strains that passed through the PIP-GEN-NIT loop. Bars represent the median MICs. *p<0.05, Kruskal-Wallis with uncorrected Dunn’s test. (F) and (G) MICs of every strain evolved in the GEN-PIP-NIT and NIT-PIP-GEN loops. Green and red circles indicate the sequenced GEN-resensitized and GEN-resistant strains that were sequenced, respectively. The x-axes indicate the drugs against which bacteria were evolved, with the MIC antibiotic listed at the top of each panel.

**Supplementary Figure 2:**
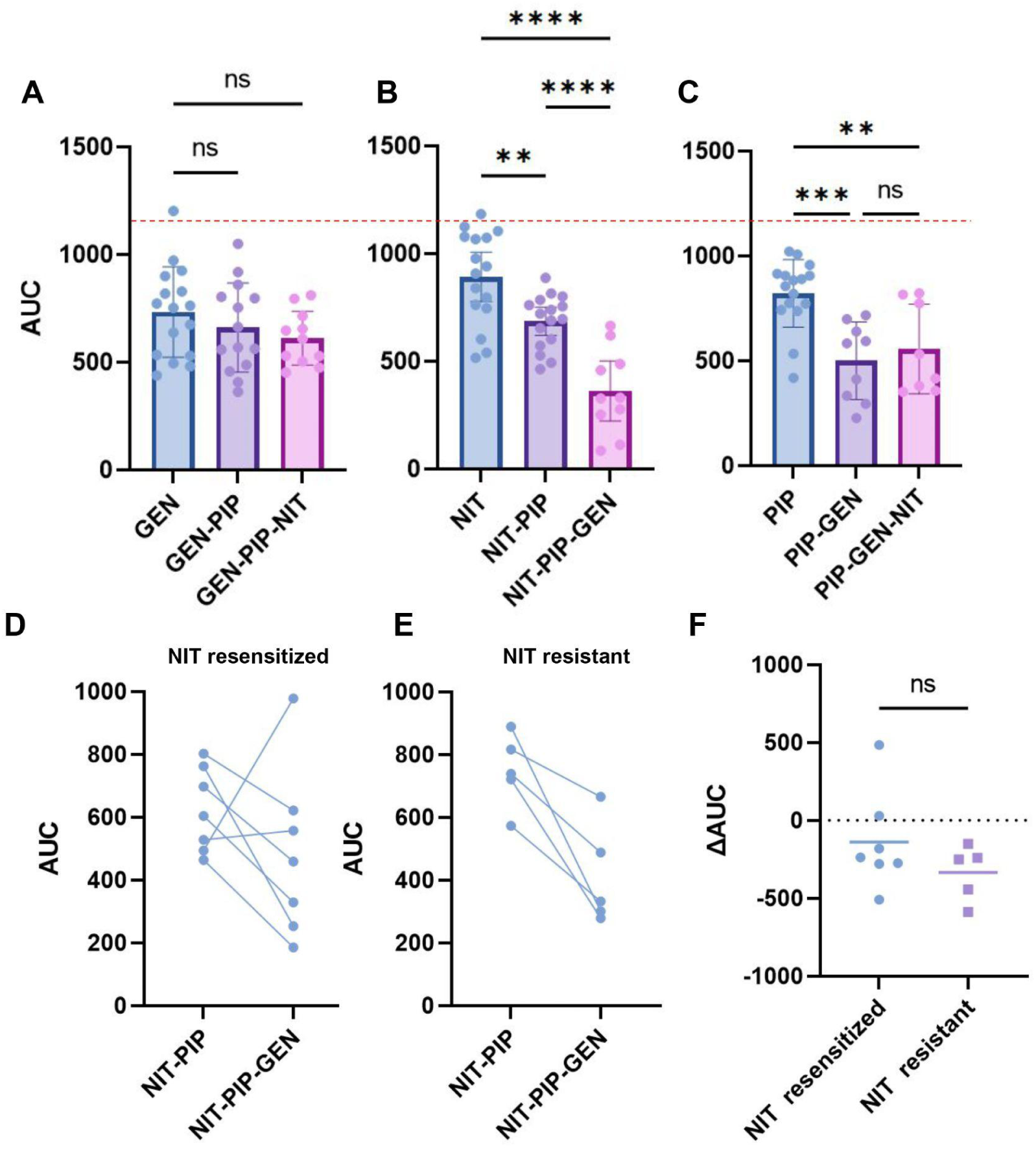
AUC data of strains passed through (A) GEN-PIP-NIT, (B) NIT-PIP-GEN, and (C) PIP-GEN-NIT loops. The red dotted line denotes the fitness of the WT strain. Bars represent the mean with 95% CI. *p<0.05, **p<0.01, ***p<0.001, ****p<0.0001, one-way ANOVA with Fisher’s LSD test. (D) and (E) AUCs of strains before and after GEN evolution for NIT-resensitized and NIT-resistant strains, respectively. The x-axes denotes the sequence of antibiotics against which the strains were evolved before measuring AUCs. (F) ΔAUC of individual strains plotted, grouped by resensitized and resistant; unpaired t test used to test significance. Means indicated by horizontal lines.

**Supplementary Figure 3:**
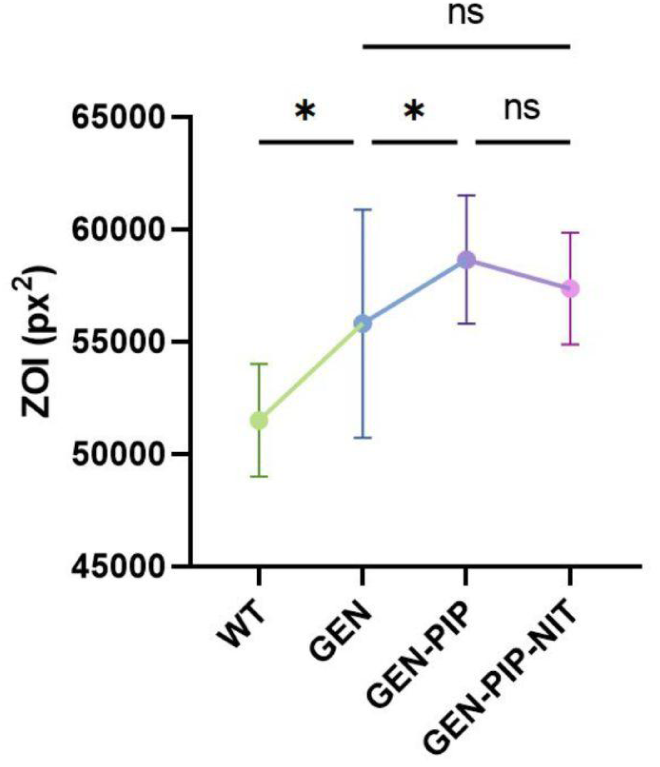
Results from the gentamicin uptake assay. GEN was incubated with bacteria (n = 3), and then centrifuged to pellet the cells. The supernatant was used to spot E. coli seeded plates (more details in Materials and Methods). The lower the GEN uptake, the more GEN remaining in the supernatant after centrifugation and hence, the larger the ZOI. Error bars represent SD. *p<0.05 one-way ANOVA with Fisher’s LSD test.

